# Liquid-liquid Phase Separation of Alpha-synuclein Increases the Structural Variability of Fibrils Formed during Amyloid Aggregation

**DOI:** 10.1101/2023.12.28.573534

**Authors:** Mantas Ziaunys, Darius Sulskis, Dominykas Veiveris, Aurimas Kopustas, Ruta Snieckute, Kamile Mikalauskaite, Andrius Sakalauskas, Marijonas Tutkus, Vytautas Smirnovas

## Abstract

Protein liquid-liquid phase separation (LLPS) is a rapidly emerging field of study on biomolecular condensate formation. In recent years, this phenomenon has been implicated in the process of amyloid fibril formation, serving as an intermediate step between the native protein transition into their aggregated state. The formation of fibrils via LLPS has been demonstrated for a number of proteins related to neurodegenerative disorders, as well as other amyloidoses. Despite the surge in amyloid-related LLPS studies, the influence of protein condensate formation on the end-point fibril characteristics is still far from fully understood. In this work, we compare alpha-synuclein aggregation under conditions, which promote or negate its LLPS and examine the differences between the formed aggregates. We show that alpha-synuclein phase separation generates a wide variety of assemblies with distinct secondary structures and morphologies. The LLPS-induced structures also possess higher levels of toxicity to cells, indicating that biomolecular condensate formation may be a critical step in the appearance of disease-related fibril variants.

## Introduction

Liquid-liquid phase separation (LLPS) of proteins and nucleic acids is a new and rapidly advancing field of study, which aims at understanding the complex process of biomolecular condensate formation ^1,2^. While this phenomenon of membranelles compartmentalization into areas of low and high protein or nucleic acid concentration was reported in various studies for more than a hundred years, it only gained significant traction over the last decade ^3,4^, when it was discovered that condensate formation plays an important role in various cellular processes ^1^. LLPS is responsible for the formation of such membranelles bodies as: promyelocytic leukemia protein bodies (involved in genome stability, programmed cell death and cell division) ^5^, stress granules (modulation of stress response, viral infection and signaling pathways) ^6^, signaling puncta (promotion of signaling outputs) ^7^, and intrinsically disordered protein condensates (onset of neurodegenerative disorders) ^3,8–11^. Currently the field lacks mechanistic information of the process governing biomolecule condensate formation and its role in diseases, which limits the search for potential treatment modalities ^12–14^. Numerous recent reports demonstrated that most of amyloid-disease related proteins are capable of undergoing LLPS ^11,15,16^, however, its influence on the aggregation process is still not fully understood ^17,18^.

The LLPS of proteins and peptides such as: amyloid-beta ^19^, Tau ^15^, TAR DNA-binding protein 43 (TDP-43) ^20^ and fused in sarcoma (FUS) ^20^ is likely an intermediate step during the transition from their native states to amyloid aggregates ^21^. However, while many reports demonstrate the formation of insoluble aggregates within the protein droplets, they do not analyse their structural characteristics. The lack of this information limits our understanding of neurodegeneration and processes involving LLPS in amyloid fibril formation such as: 1) its influence on the secondary structure ^22^ and morphology of the resulting aggregates ^23^, 2) their self-replication properties ^24^, 3) toxicities ^15^, and 4) susceptibility to anti-amyloid compounds ^25^. Addressing the questions related to LLPS in amyloid fibril formation is important for advancing our understanding of neurodegenerative diseases, unravelling fundamental biophysical processes, and informing drug development efforts aimed at combating these debilitating conditions.

Aggregation of the intrinsically disordered protein – alpha-synuclein (α-syn) ^26^ into amyloid fibrils is associated with the second most common incurable neurodegenerative disorder – Parkinson’s disease ^27^. After decades of intensive research, there are still no effective cures or treatment modalities available for both this, as well as many other amyloid-related disorders ^28,29^. The lack of mechanistic understanding of different steps in the aggregation process of α-syn and other disease-related proteins pose a significant obstacle in the search for effective disease-altering drugs ^30,31^. The newly discovered LLPS is potentially an intermediate state between native and fibrillar forms of amyloidogenic proteins that plays a crucial role in determining the resulting aggregate structure, self-replication potential and cytotoxicity. This hypothesis is based on the fact that α-syn amyloid aggregation is significantly influenced by even minor alterations in environmental conditions, such as pH or ionic strength ^32–35^. This includes protein concentration, which becomes considerably larger within droplets ^36,37^, as well as the appearance of a liquid-liquid interface, possibly altering the fibril formation process ^38^.

In this work, we design a novel approach for LLPS studies *in situ* and utilise it to examine how LLPS influences the aggregation kinetics, resulting fibril secondary structure, morphology, cytotoxicity and self-replication potential of α-syn. Next, we compare these characteristics against fibrils prepared under non-LLPS conditions. The results of the study demonstrate that α-syn incubation under LLPS conditions leads to a wide variety of structures, ranging from small and unstable amorphous aggregates to several types of fibrils with varying morphological characteristics. These findings suggest that LLPS may be a critical process in the formation of disease-related fibrils, which differ significantly from ones generated during commonly used *in vitro* approaches.

## Materials and Methods

### Cloning eGFP-alpha-synuclein

The EGFP-alpha-synuclein gene was amplified from Addgene plasmid # 40822 (EGFP-alpha-synuclein-WT was a gift from David Rubinsztein (http://n2t.net/addgene:40822; RRID:Addgene_40822). The forward (5’-GCCCATGGTGAGCAAGGGCGAGGAG-3’) and reverse (5’-GTCGCCATCGAAGCTTGAGCTCGAGATCTG-3’) primers were used to insert *NcoI* and *XhoI* sites, followed by traditional restriction enzyme cloning into the pET28B(+) vector ^39^.

### Alpha-synuclein purification

Alpha-synuclein (α-syn) and eGFP-α-syn was purified based on a similar version of a previously described protocol ^40^. In short, the pRK172 vector with human alpha-synuclein was expressed in *E.coli* BL-21StarTM (DE3) cells in ZYM-5052 medium (containing 100 µg/ml ampicillin) at 37 °C. After overnight synthesis, cells were pelleted, then homogenised and lysed in 20 mM Tris-HCl (pH 8.0) buffer solution, containing 0.5 M NaCl, 1 mM EDTA and 1 mM PSMF. The collected homogenate was heated to 70°C in a water bath and incubated with stirring for 10 minutes. The formed precipitates were pelleted by centrifugation at 12 000 x g for 15 minutes. The supernatant was filtered through a 0.22 µm pore size syringe filter and cooled down to 4°C. Ammonium sulphate was added to the solution to a saturation of 42% and stirred at 300 RPM for 30 min. Precipitates formed during this procedure were pelleted by centrifugation at 12 000 x g for 15 minutes. The pellets were then dissolved in a 20 mM Tris-HCl (pH 8.0) buffer solution, containing 0.5 mM DTT and filtered using a 0.22 µm pore size syringe filter. The solution was loaded onto a DEAE sepharose sorbent for ion exchange chromatography. Unbound proteins were washed using 5 column volumes of the 20 mM Tris-HCl (pH 8.0), 0.5 mM DTT buffer solution. α-syn was eluted using a 20 mM Tris-HCl (pH 8.0), 0.5 mM DTT, 200 mM NaCl buffer solution. Collected fractions containing α-syn were concentrated and further purified using a HiLoad 26/600 size exclusion chromatography (SEC) column (packed with Superdex 75 resin) where α-syn was eluted at 2.5 ml/min using PBS. After SEC, the protein solutions were frozen at −20°C. To avoid batch-to-batch variability, protein fractions from multiple SEC were thawed at room temperature, combined, concentrated and filtered through 0.22 µm pore-size syringe filters yielding a final protein concentration of 600 µM. The protein solution was distributed into fractions of 500 µL and stored at −20°C. The protein stock solution was used within 3 months of purification.

### Alpha-synuclein liquid-liquid phase separation and aggregation

For LLPS-induced aggregation, the α-syn stock solution was combined with a 30% polyethylene glycol (PEG, 20 kDa) PBS solution, containing 150 µM thioflavin-T (ThT) to yield a final protein concentration of 200 µM, 20% PEG and 100 µM ThT. The reaction solution was then distributed to 24 wells in the center of Greiner Bio-one 96-well non-binding surface plates (200 µL in each well, sample placement is shown as Supplementary Figure S1). The plates were then incubated under quiescent conditions at 37°C for 96 hours in a ClarioStar Plus plate reader. Sample optical density at 600 nm and ThT fluorescence intensity (440 nm excitation, 480 nm emission) were simultaneously scanned every 10 minutes. After the 96-hour incubation period, the plates were cooled down to room temperature and samples were recovered from each well for further analysis.

For positive and negative control aggregation kinetics, the reaction solutions were prepared without PEG. In the case of the negative control, the reaction solution was distributed and incubated as during LLPS-induced aggregation. For the positive control, each well contained an additional 3 mm glass bead, and the plate was subjected to constant 600 RPM orbital agitation. In this case, sample optical density was not tracked due to the signal being affected by the presence of the 3 mm glass beads.

### Aggregate dissociation

50 µL aliquots of samples from the aggregation kinetic assays were pooled together (24 repeats, total final volume was 1.2 mL) in non-binding 2 mL test-tubes and centrifuged at 14 000 x g for 20 min at 22°C. The supernatants were removed (1.1 mL) and the aggregate pellets were resuspended into PBS solutions (final volume – 1.2 mL). The centrifugation and resuspension steps were repeated four times. After the final resuspension procedure, the sample optical densities at 600 nm (OD_600_) were scanned using a Shimadzu UV-1800 spectrophotometer in a 3 mm pathlength cuvette. The samples were then separated into equal fractions for denaturant-induced and cold-induced aggregate dissociation assays.

For cold-induced dissociation ^41^, the samples were incubated at 4°C under quiescent conditions. The optical density measurements were performed every 24 hours, after brief sample mixing by pipetting for 5 seconds. Prior to scanning the OD_600_ values, the aliquots (100 µL) were placed in the cuvettes and incubated at 22°C for 5 minutes. For denaturant-induced dissociation, aliquots of the samples (50 µL) were combined with 50 µL PBS solutions, containing different concentrations of guanidinium hydrochloride (GuHCl, from 0 M to 2 M in 0.2 M increments). The final solutions contained 100 µM α-syn and GuHCl concentrations from 0 M to 1 M in 0.1 M increments. The solutions were then incubated at 22°C for 1 hour, mixed by pipetting for 5 seconds, after which their optical densities were scanned.

### Self-replication

To compare the self-replication properties of LLPS-induced aggregates with the positive control ones, both samples were subjected to the same resuspension and cold-incubation procedures. They were then combined with the initial α-syn reaction solutions without PEG (20% aggregate solution) and incubated under quiescent conditions at 37°C in 96-well non-binding plates. The self-replication reactions were tracked by monitoring the sample bound-ThT fluorescence intensity in a ClarioStar plus plate reader.

### Atomic force microscopy (AFM)

Aliquots of each sample (50 µL) were diluted 10 times with PBS and sonicated for 10 seconds using a Bandelin Sonopuls ultrasonic homogenizer, equipped with a MS-72 sonication tip (20% intensity). 30 µL aliquots of each sample were placed on APTES-modified mica and washed using 3 mL of Milli-Q H_2_O. The AFM image acquisition was done as previously described ^42^. In short, 51×5 µm images of 10241×1024 pixel resolution were acquired for each sample using a Bruker Dimension Icon atomic force microscope, operating in tapping-mode. The images were flattened and analysed using Gwyddion 2.62 software.

Particle count was performed by selecting three separately acquired AFM images and tracing all particles present in a selected area (1×1 µm for initial aggregate samples, 2.51×2.5 µm for samples after cold-dissociation and 51×5 µm for samples after reseeding). Individual aggregate particles were all spatially separated AFM image elements with >1 nm cross-sectional height. Particles, which had an elongated structure (particle length was more than three times larger than its width) were considered as fibrils.

For elongated fibril analysis, the samples were not sonicated prior to their placement on APTES-modified mica. 1 µm length fibril images were cut from the large-scale AFM images and exported as raw height data. The raw data was fit using a Gaussian function (Origin Software) and the fit results were used to determine the fibril’s cross-sectional height and width pixel-by-pixel along the axis of each aggregate.

### Fourier-transform infrared (FTIR) spectroscopy

For time-course measurements of α-syn secondary structure changes during incubation under 0% and 20% PEG conditions, the protein had to be exchanged from PBS solution to an equivalent heavy water solution. D_2_O PBS was prepared by replacing hydrogen-containing PBS components with their deuterated variants. The pD of the solution was set to mimic the pH of PBS using a previously described conversion formula ^43^. pD corrections were done by using DCl and NaOD. α-syn was exchanged into deuterated PBS using a 10 kDa cut-off protein concentrator with 5 rounds of dilution and concentration. The 30% PEG stock solution was prepared by dissolving 20 kDa PEG in deuterated PBS, after which the pD value was corrected. Further reaction solution preparation was done as in the previously described regular PBS LLPS and aggregation assays.

To prevent H_2_O/D_2_O exchange during the long incubation period, the samples were placed in 2 mL non-binding test-tubes (1.5 mL volume each) and sealed using parafilm. The test-tubes were incubated under quiescent conditions under a constant 37°C temperature. For measurements at each selected time-point, the samples were gently rotated 5 times to resuspend any possible aggregates, after which 30 µL aliquots were taken and the samples were resealed. The aliquots were cooled down to room temperature and scanned using a Bruker Invenio S FTIR spectrometer. The scanning procedure was performed as described previously ^44^. Data processing was done using GRAMS software. D_2_O, water vapour and PEG spectra were subtracted from the sample spectra, which were then baseline corrected and normalized to the same area between 1700 cm^−1^ and 1600 cm^−1^.

### Aggregate concentration determination

Samples were centrifuged at 14 000 x g for 15 minutes, after which the supernatant was removed and replaced with an identical volume of PBS solution, this procedure was repeated three times in order to remove any residual, non-aggregated protein. After the final resuspension, the solution was combined with a 6 M GuHCl PBS solution (1:4 ratio) and incubated at room temperature for 1 hour. After this procedure, the sample absorbance at 280 nm was scanned using a Shimadzu UV-1800 spectrophotometer. The obtained absorbance values were used to determine the aggregated protein concentrations in the initial samples (ε_280_=5960 M^−1^cm^−1^).

### Fluorescence microscopy

For fluorescence imaging experiments, the formed droplets of α-syn with 1% of eGFP-α-syn and/or α-syn with 100 µM ThT were immobilized on a bare glass surface. This was done by slowly adding 100 µL of droplet suspension onto a cleaned uncoated glass coverslip (Menzel Coverslip 241×60mm #1.5 (0.16 – 0.19 mm), Thermo Scientific, cat. no. 17244914) and no subsequent washing steps were performed. To minimize the possibility of droplet collapse during their physical manipulation, wide-orifice tips (Finntip 250 Wide, Thermo Scientific, cat. no. 9405020) were used without any intense pipetting of the droplet suspension. Single-molecule fluorescence microscopy imaging of such samples was performed using a home-built super-resolution microscope – the miEye ^45,46^. To visualize the individual glass surface-attached α-syn droplets, TIRF microscopy was employed ^47^, whereas imaging of larger α-syn formations, such as droplet aggregates, was conducted under HILO illumination. Lasers of 405 and 488 nm wavelengths were used to excite α-syn-ThT and α-syn-eGFP, respectively, and the resulting fluorescence was collected through a 525/45 nm band-pass filter. In all cases, the exposure time of an industrial CMOS camera (Alvium 1800 C-511m, Allied Vision Technologies) was set to 100 ms.

### MTT and LDH assays

Before conducting the MTT and LDH assays, all sample protein concentration required to be equal. Monomeric α-syn and non-LLPS aggregates were diluted to 20 µM using PBS. LLPS aggregate samples were centrifuged at 14 000 x g for 15 min and the aggregate pellets were resuspended into a smaller volume of PBS, resulting in a final protein concentration of 20 µM.

Cell culture used for MTT and LDH assays (SH-SY5Y human neuroblastoma cells) were obtained from the American Type Culture Collection (ATCC, Manassas, VA, USA). The cells were grown in Dulbecco’s Modified Eagle Medium (DMEM), supplemented with 10% Fetal Bovine Serum (FBS), 1% Penicillin– Streptomycin (10 000 U/mL), at 37 °C in a humidifier, 5% CO_2_ atmosphere in a CO_2_ incubator.

For the MTT assay, SH-SY5Y cells were seeded in a 96-well plate (∼15 000 cells/well) and were incubated overnight. After incubation, the medium was changed to one containing different forms of α-syn (monomers or fibrils). Since all samples were prepared in PBS, the medium with corresponding PBS amount was used as a control. After 48 h of incubation, 10 µM of 3-(4,5-dimethylthiazol-2-yl)-2,5-diphenyltet-tetrazolium bromide (MTT) reagent (12.1 mM in PBS) was added to each well, followed by 2 h of incubation. To dissolve formazan crystals, 100 µL of 10% SDS with 0.01 N HCl solution was added to each well. After 2 h, the absorbance was measured at 570 nm and 690 nm (as reference wavelength) using a ClarioStar Plus plate reader.

The LDH assay was performed as previously described ^48^. SH-SY5Y cells were seeded in a 96-well plate (15,000 cells/well) and were incubated overnight. After incubation the medium inside the wells was aspirated, 100 µL/well of Advanced DMEM was added. Then, each sample containing α-syn was diluted in Advanced DMEM. Prepared samples were added to the wells to reach 200 µL/well of total medium volume, which resulted in the final 5 µM α-syn sample concentration. After 24 h of incubation, 100 µL of the medium from each well was aspirated, centrifugated, and transferred into a TPP 96-well tissue culture test plate. Freshly prepared LDH reagent was added and incubated for 30 min at room temperature. The absorbance was measured at 492 nm and 600 nm (as reference wavelength) using a ClarioStar Plus plate reader.

## Results

LLPS of α-syn can be induced with the addition of a crowding agent (PEG) to a high protein concentration sample (200 µM final concentration). The hypothesis is that LLPS would alter the process of fibril formation and affect their structural characteristics in a way that they become different from aggregates generated without LLPS. To track the formation of protein droplets, as well as α-syn aggregates, we employed time course sample optical density measurements. Simultaneously, the appearance of amyloid fibrils was observed by using an amyloid-specific dye –thioflavin-T (ThT), which selectively binds to their beta-sheet grooves ^49^. To test the aforementioned hypothesis, α-syn was subjected to three different conditions: 1) incubated under quiescent conditions with no crowding agent (negative control), 2) incubated with constant 600 RPM agitation (positive control), and 3) incubated under quiescent conditions with 20% PEG (LLPS-inducing conditions). The negative control showed no detectable changes in both the fluorescence intensity of ThT and sample optical density (Figure 1A), indicating that no droplet formation or amyloid aggregation took place during the 96-hour period. The positive control showed the formation of amyloid fibrils within the initial 24 – 48 hours (Figure 1C). The observed large variations in raw fluorescence intensity values were due to previously reported high quantum-yield ThT binding modes, originating from subtle morphological and structural aggregate differences ^33^. The LLPS-inducing condition experiment showed changes in sample fluorescence intensity and optical density that did not follow an identical pattern (Figure 1B). The ThT fluorescence intensity had a minor increase during the initial hours of incubation, followed by a lag phase and a substantially larger change past the 24-hour point. Optical density had a comparable initial growth; however, it did not possess a similar increase after the 24-hour mark and continued to gradually change during the entire monitoring period.

**Figure 1.**
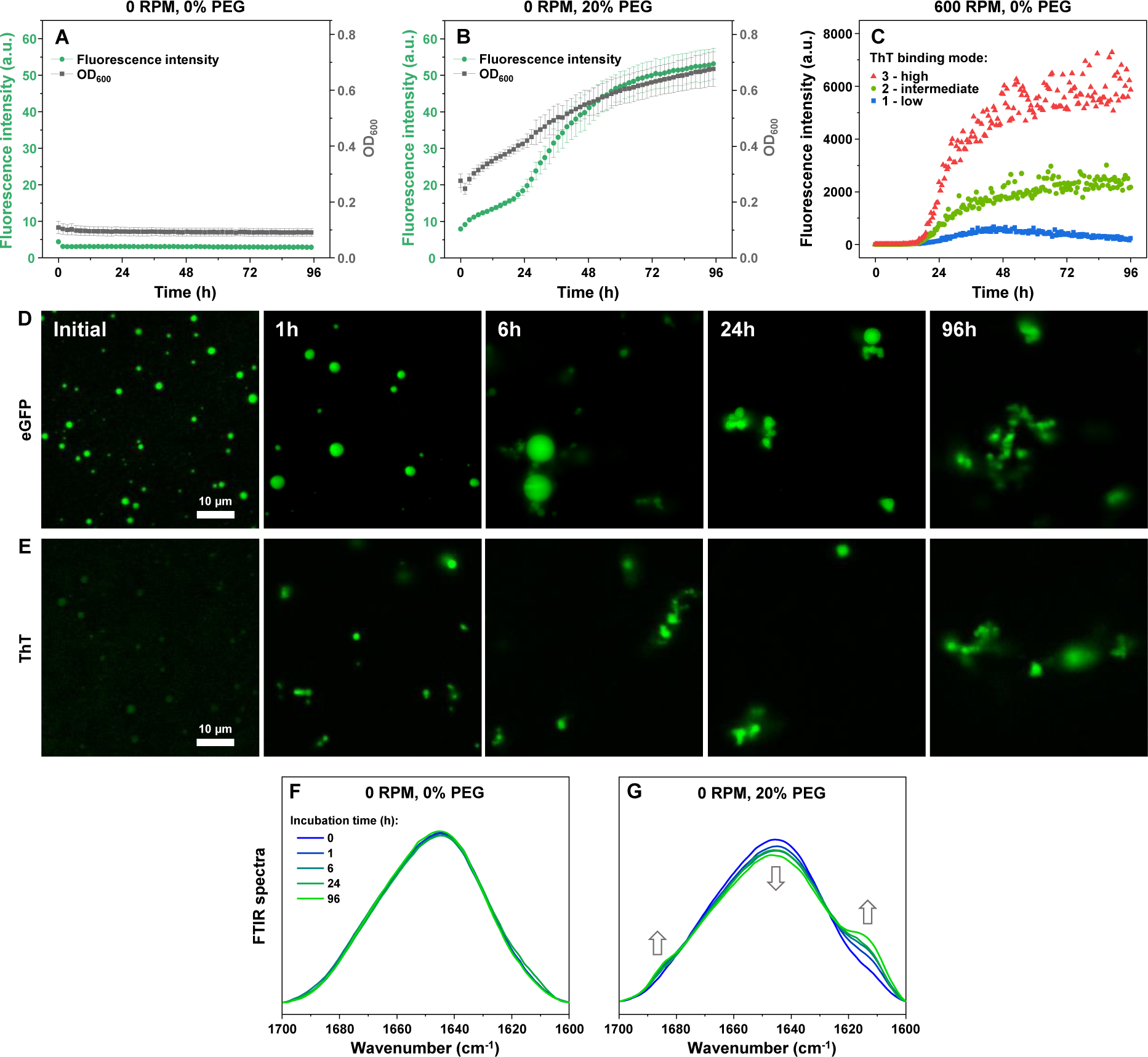
Alpha-synuclein liquid-liquid phase separation and aggregation. Thioflavin-T fluorescence intensity and sample optical density changes of alpha-synuclein under quiescent conditions without PEG (A, negative control), with 20% PEG (B, LLPS-inducing conditions) and under 600 RPM agitation without PEG (C, positive control). Fluorescence microscopy images of eGFP-tagged alpha-synuclein (D) and ThT-dyed samples (E). For each respective set of images, the intensity scale minimum and maximum values are identical. Changes in FTIR spectra of samples incubated in the absence (F) or presence of 20% PEG (G) under quiescent conditions.

Fluorescence microscopy was used to examine the appearance of α-syn droplets, alterations in their size and subsequent aggregate formation. For this task, protein samples with either 1% eGFP-tagged α-syn or 100 µM ThT were incubated under LLPS-inducing conditions. Aliquots of both samples were taken multiple times during the incubation period for analysis. The initial samples were immobilized on a bare glass surface 10 minutes after mixing the protein with PEG and immediately imaged. Despite the short time period, the samples already contained protein droplets (Figure 1D), similar to a previous report of rapid α-syn LLPS ^50^. Interestingly, the droplets were also capable of binding ThT molecules in a mode, which facilitated their fluorescence to overcome the background level (Figure 1E). This suggests that the initial droplets already had a small number of ThT-binding structures. After 1 hour of incubation, there were no discernible differences in the case of the eGFP-α-syn samples, while the ThT sample droplets were noticeably brighter than in the initial samples, indicating an increase in the concentration of ThT-positive structures. After 6 hours, the samples contained small amounts of enlarged droplets, as well as fibrillar and amorphous structures, visible in both the eGFP and ThT samples. Further incubation led to the amorphous, ThT-positive structures becoming the dominant population. In the case of the negative control samples, there were no discernible droplet or aggregate formations visible throughout the entire incubation time-period (Supplementary Figure S2). Additional larger scale fluorescence microscopy images of both sample types are shown as Supplementary Figures S3 and S4.

To gain insight into the secondary structure changes of α-syn during LLPS, the protein samples were examined at identical time-points as in the fluorescence microscopy assay using Fourier-transform infrared spectroscopy (FTIR) and compared against the negative control. The initial samples had highly similar FTIR spectra (Figure 1F, G), with slight differences in the ∼1650 cm^−1^ (random-coil) and ∼1620 cm^−1^ (parallel beta-sheets) positions ^51^. Further incubation led to accentuated distinctions between the spectra (Figure 1E, G), with LLPS conditions resulting in a reduction in the band associated with random-coils and the formation of anti-parallel (∼1695 cm^−1^) and parallel (∼1620 cm^−1^) beta-sheets ^51^. Despite these alterations in the FTIR spectra, the 96-hour spectrum (Figure 1G) suggested the presence of either large quantities of non-aggregated α-syn or amorphous, randomly organized structures. Oppositely, the negative control conditions did not result in any significant alterations to the FTIR spectra throughout the entire incubation period (Figure 1F).

The presence of such an abundance of amorphous structures posed an issue for further structural characterization of amyloid fibrils, formed during LLPS-promoting conditions. To solve this problem, two different approaches were applied. First, we induced amorphous aggregate disassembly while retaining the α-syn fibrils using sample denaturation by low concentrations of guanidinium hydrochloride (GuHCl)^52^. The LLPS and non-LLPS (positive control) condition aggregates were resuspended into a range of GuHCl concentrations (Figure 2A), and the effect of denaturation was monitored by scanning sample optical densities (OD_600_). In the case of LLPS aggregates, there was a sharp decrease in the sample optical density up to 0.6 M GuHCl, followed by a discontinuity in the trend. Further increase in GuHCl concentration had a considerably lower effect on sample OD_600_ values. Non-LLPS aggregates were significantly less susceptible to denaturation up to 0.6 M GuHCl, followed by a sharper reduction in OD_600_. Interestingly, the trend discontinuities both appeared at the same GuHCl concentration, suggesting that LLPS and non-LLPS samples contained different concentrations of similarly unstable aggregates, with them being the dominant population in LLPS samples.

**Figure 2.**
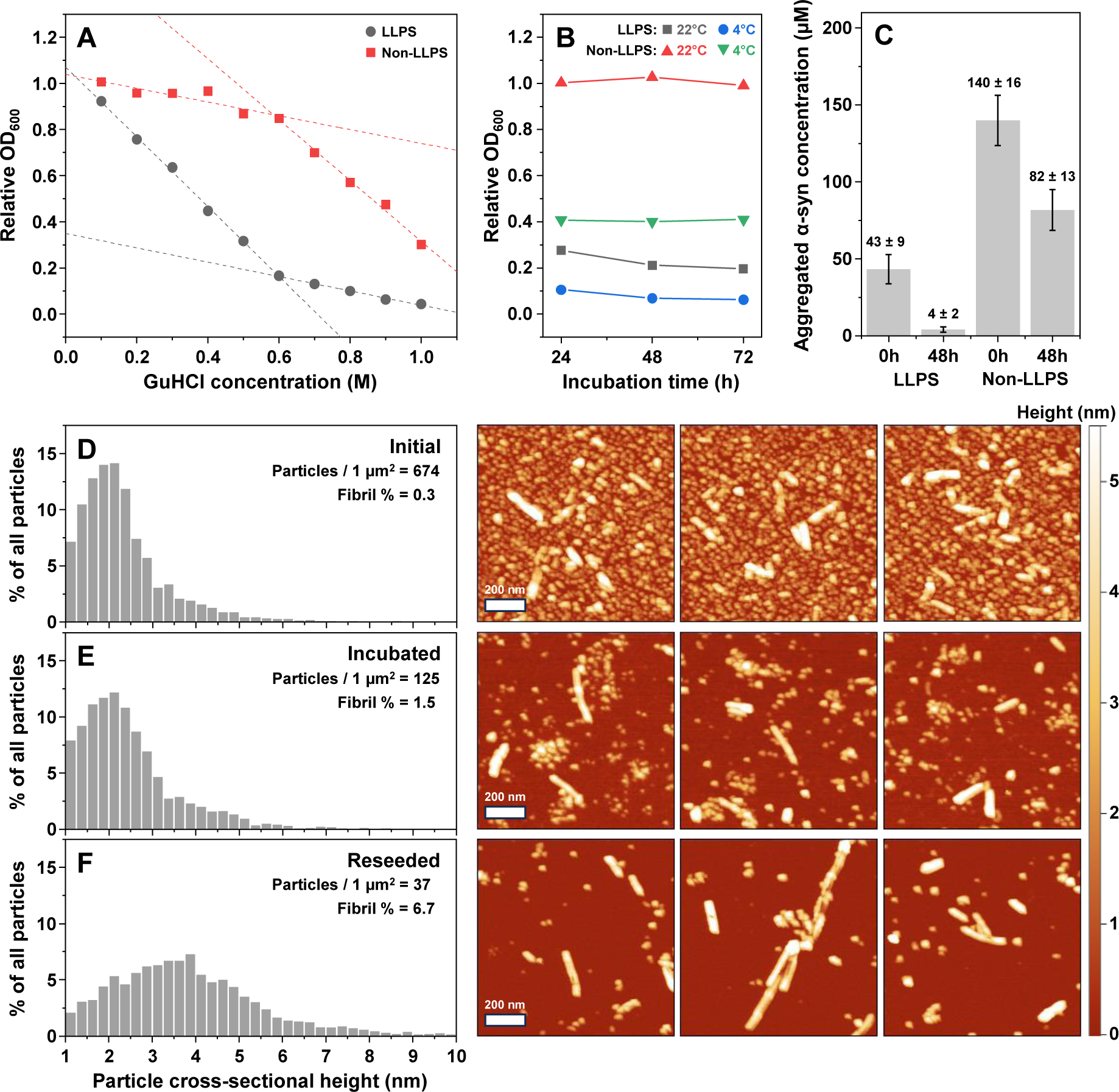
Isolation of stable α-syn fibrils from initial aggregate mixtures. LLPS and non-LLPS sample relative optical density at 600 nm (OD_600_) dependence on guanidine hydrochloride (GuHCl) concentration (A) and incubation at different temperatures (B). Aggregated α-syn concentration in LLPS and non-LLPS samples before and after cold incubation (C). Particle cross-sectional height distribution and atomic force microscopy image examples of initial (D), cold-incubated (E) and reseeded (F) LLPS samples.

Second, we attempted to destabilise α-syn aggregates under low temperature conditions ^41^. LLPS and non-LLPS aggregates were resuspended into PBS without the crowding agent and incubated at 22°C and 4°C for multiple days. As expected, the OD_600_ of non-LLPS samples at room temperature remained unchanged during the entire incubation period (Figure 2B). In the case of LLPS aggregates, the mere act of resuspension into PBS and incubation at 22°C led to a large reduction of sample OD_600_. Cold incubation caused a high level of disassembly for both LLPS and non-LLPS samples (0.1 and 0.4 relative OD_600_ respectively). Interestingly, incubation under 4°C conditions resulted in similar relative OD_600_ values as resuspension of the aggregates in 1.0 M of GuHCl (Figure 2A, B). Measuring the concentrations of non-aggregated α-syn before and after cold incubation (Figure 2C) revealed that LLPS only caused (43 ± 9) µM of the protein to form insoluble structures. Incubation at 4°C reduced this number to (4 ± 2) µM, indicating that only a tiny fraction of structures in LLPS samples had considerable stability. Oppositely, non-LLPS conditions caused the aggregation of the majority of α-syn (140 ± 16) µM and cold incubation only reduced the concentration of insoluble structures to (82 ± 13) µM. Considering that cold incubation resulted in a comparable level of disassembly as 1.0 M GuHCl and circumvented the need for aggregate separation from the denaturant, this method was used as a means of stable α-syn structure isolation from the prepared samples.

To determine the distribution of round, amorphous structures and fibrillar aggregates during the cold incubation and self-replication procedures we employed atomic force microscopy (AFM). After the initial resuspension of LLPS condition aggregates into PBS without the crowding agent, AFM images displayed a massive amount of different size round aggregates, covering almost the entire surface of the mica (675 particles/µm^2^), with a few fibrillar structures observed in-between (Figure 2D). Based on the particle count, only 0.3% of them possessed an elongated structure, while the majority of round structures had a cross-sectional height of ∼2 nm. After cold incubation, the particle count was reduced to 125/µm^2^ and the relative abundance of fibrillar structures increased to 1.5% (Figure 2E). After reseeding, the particle count was even further reduced to just 37/µm^2^ and the percentage that had fibrillar shapes was 6.7% (Figure 2F). The particle cross-sectional height distribution peak was also shifted from 2 nm to 4 nm. These results display the effectiveness of utilizing cold incubation, in combination with reseeding, to massively reduce the number of round and amorphous structures, while retaining and increasing the concentration of fibrils.

To compare the LLPS aggregates with the positive control fibrils, the entire process of resuspension, cold incubation and reseeding was performed with both sample types. It was observed, however, that LLPS aggregates were not nearly as effective at self-replication under conditions without crowding agents, as compared to non-LLPS fibrils (Supplementary Figure S5). AFM images were acquired for reseeded LLPS and non-LLPS samples with the exclusion of the sonication step, which was used for particle analysis. Large scale AFM images of both aggregate types are available as Supplementary Figures S6 and S7. Fibrils from each image were then analysed by tracing lines perpendicular to the fibril axes in a pixel-by-pixel basis and acquiring information about the morphology of the entire lengths of the different fibrils.

Conditions, which facilitated LLPS, resulted in a variety of fibrillar structures (Figure 3A), a phenomenon previously observed for α-syn aggregates under a number of specific environmental conditions ^33–35^. Oppositely, the non-LLPS aggregates shared a highly similar morphology, with little to no variation in between separate AFM images (Figure 3B, Supplementary Figure S6). Out of multiple LLPS condition AFM images, five unique fibril types were detected and analysed (Figure 3C-G). Four of the aggregate types contained clearly visible periodicity patterns (as seen in the fibril height heatmaps (Figure 3C-G). The periodicities were all different and ranged from as low as (58 ± 8) nm in the case of Type III aggregates, to (328 ± 27) nm – Type II. Type V fibrils contained alternating regions of different cross-sectional heights; however, no repeating periodicity was detected (Figure 3G).

**Figure 3.**
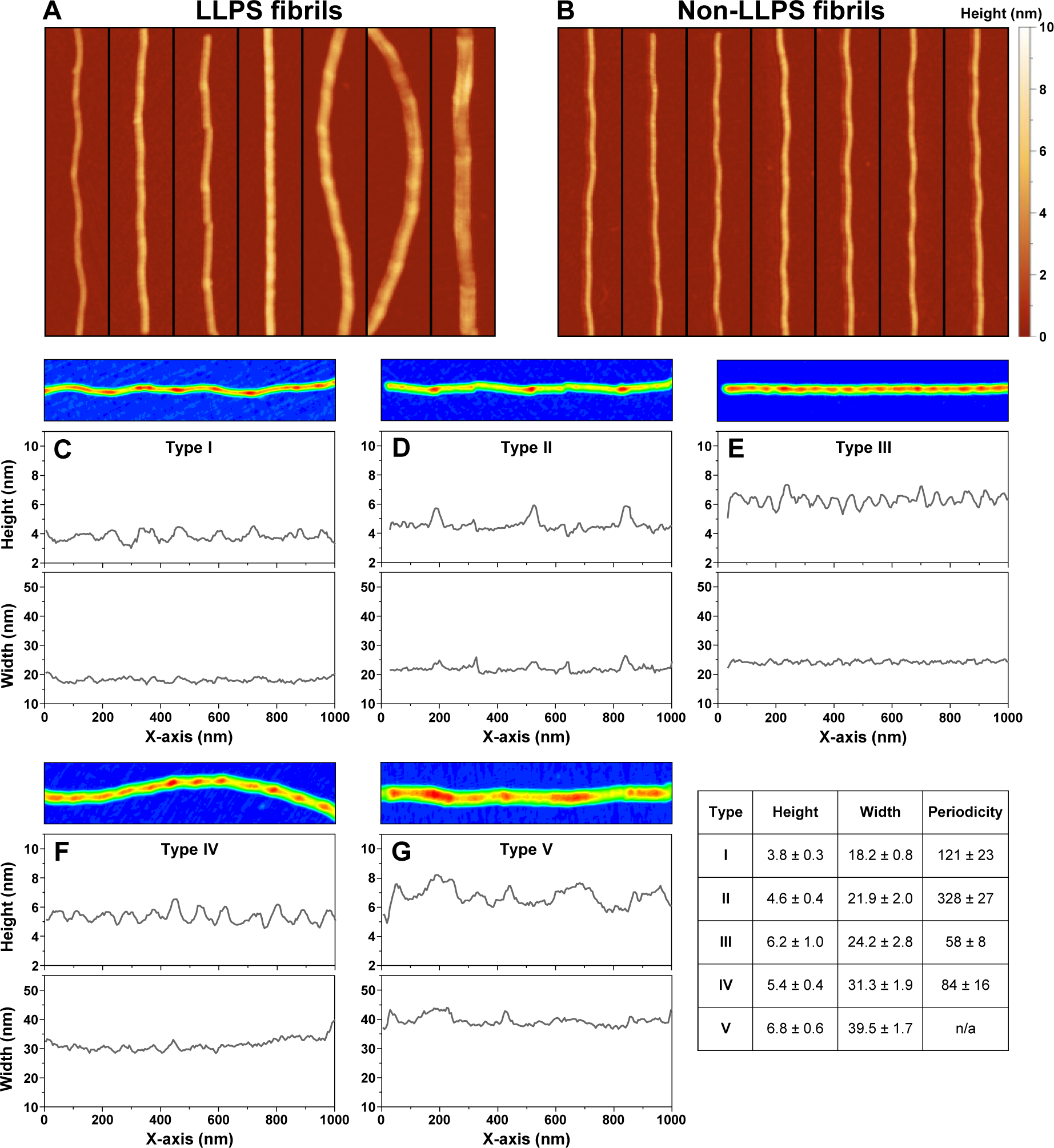
Morphology of α-syn fibrils in LLPS and non-LLPS samples. Examples of LLPS (A) and non-LLPS (B) fibrils, found among the several larger scale AFM images. Different fibril AFM image heatmaps and their respective height and width values along the axes of Type I (C), II (D), III (E), IV (F) and V (G) fibrils. Table insert shows the height, width, and periodicity of each fibril type. Height and width values were determined along the entire axes of each fibril type as described in the Materials and Methods section. Periodicity is the average distance between fibril height local maximum positions (n>4 for all types, error values are one standard deviation).

In addition to distinct periodicities, the five fibril types also had varying average height and width values (Figure 3 Table insert). Type I fibrils had the lowest average cross-sectional height and width ((3.8 ± 0.3) nm and (18.2 ± 0.8) nm respectively), followed close-second by Type II aggregates ((4.6 ± 0.4 nm and 21.9 ± 2.0) nm respectively). The largest average cross-sectional height and width values were observed in the case of Type V fibrils ((6.8 ± 0.6) nm and (39.5 ± 1.7) nm respectively). The remaining two aggregate types possessed intermediate values, which did not follow a similar correlation between height and width. While Type III fibrils did not have a significantly different cross-sectional height than Type V, their width values were almost two-fold lower. Oppositely, Type IV aggregates were similar in height to Type II, while possessing a substantially larger width. Interestingly, the structures observed in non-LLPS AFM images (Figure 3A) all shared identical morphological properties as LLPS condition Type I fibrils (Figure 3C). These results indicate that this specific fibril type is capable of assembling during both LLPS-promoting conditions, as well as without the formation of droplets.

The initial, cold-incubated and reseeded aggregate samples were also examined by Fourier-transform infrared spectroscopy to determine differences in their secondary structure. The resulting spectra were deconvoluted to calculate the distribution of beta-sheet, random-coil and turn/loop motifs in the protein secondary structure. Comparing the initial LLPS and non-LLPS sample FTIR spectra (Figure 4A, D), the most notable aspect is the difference in the quantity of random-coil and beta-sheet content present in the aggregate structure (Figure 4G, H). In the case of LLPS, the aggregates possessed significantly more random-coil motifs (∼45%, as opposed to 5%), as well as less parallel beta-sheets of both weaker and stronger hydrogen bonding variant (13% and 12%, as opposed to 25% and 35% respectively). The LLPS sample spectra also had a small peak at 1695 cm^−1^, associated with anti-parallel beta-sheet hydrogen bonding. Finally, the initial LLPS sample spectra displayed a slightly lower content of turn/loop motifs (28% as opposed to 32%).

**Figure 4.**
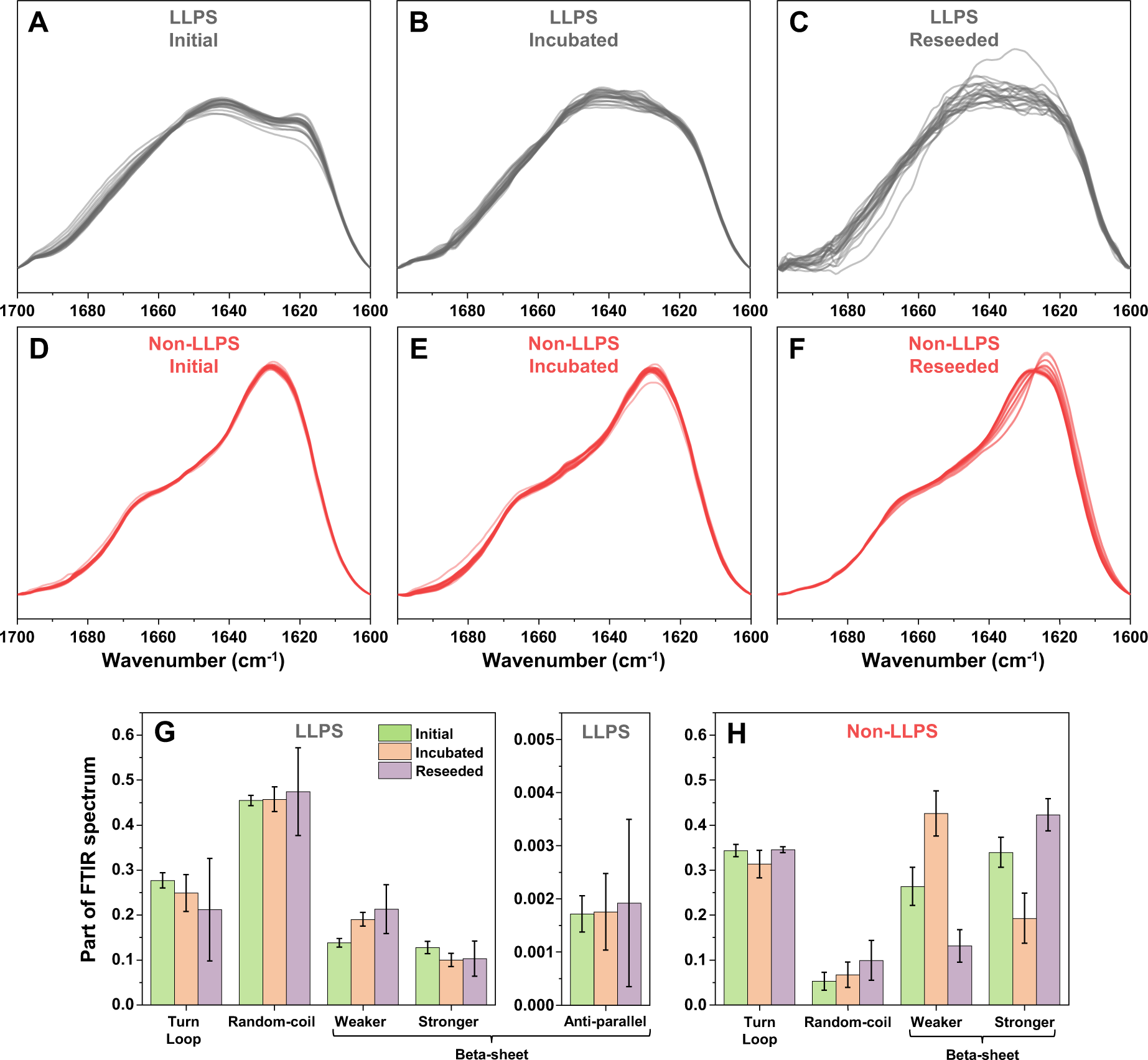
Initial, cold-incubated and reseeded LLPS and non-LLPS aggregate secondary structure distributions. FTIR spectra of initial (A, D), cold-incubated (B, E), reseeded (C, F) LLPS and non-LLPS aggregates respectively. Turn/loop, random-coil and beta-sheet motif distribution of LLPS (G) and non-LLPS (H) samples, based on FTIR spectra deconvolution (n=24, errors bars are for one standard deviation).

When the samples were cold-incubated, the only significant change for LLPS condition aggregates was an increase in the content of weaker hydrogen bond strength parallel beta-sheets (Figure 4B, G). In the case of non-LLPS, a similar change was also accompanied by a significant reduction in the content of stronger hydrogen bonding beta-sheets (Figure 4E, H). For both aggregate types, the cold-incubation did not have a notable effect on the random-coil motif, suggesting that either the unstable aggregate types had similar random-coil regions as the other assemblies, or the dissociation also affected some of the fibrillar structures. Based on the change in non-LLPS FTIR spectra and the previously observed cold-induced increase in fibril content of LLPS samples, the later possibility appears more likely.

LLPS aggregate reseeding did not cause any significant changes to the secondary structure motif distribution, however, it greatly increased the stochasticity of the spectra (Figure 4C, G). During the FTIR analysis, it was also observed that the reseeded LLPS samples contained less aggregates than the non-LLPS samples, which contributed to the higher level of noise in their spectra. These results, in combination with the previously shown reseeding kinetics, further support the hypothesis that LLPS aggregate self-replication is far less effective under non-LLPS conditions. In the case of non-LLPS samples, the reseeded aggregate spectra contained an even higher relative amount of stronger hydrogen bonding beta-sheets than the initial samples (Figure 4F, H). Some of the non-LLPS sample FTIR spectra also had a shifted main maximum position, which was the result of the aforementioned change between weaker and stronger beta-sheet content.

Due to the large differences in both the secondary structure, as well as morphology of LLPS and non-LLPS samples, it was investigated whether they also possessed distinct effects on SH-SY5Y human neuroblastoma cells. In the MTT assay, it was observed that non-LLPS and both types of LLPS aggregates (before and after 48h cold incubation) reduced cell metabolic activity (Figure 5A), however, there were no significant differences between them (n=3, p<0.05). An examination of the aggregate cytotoxicity revealed that the 48h cold incubation sample possessed the most potent effect on the cell membrane, which was significantly higher than non-LLPS aggregates (Figure 5B). The 0h incubation sample results had a relatively high error and no statistically significant differences from non-LLPS and the 48h incubation sample could be determined.

**Figure 5.**
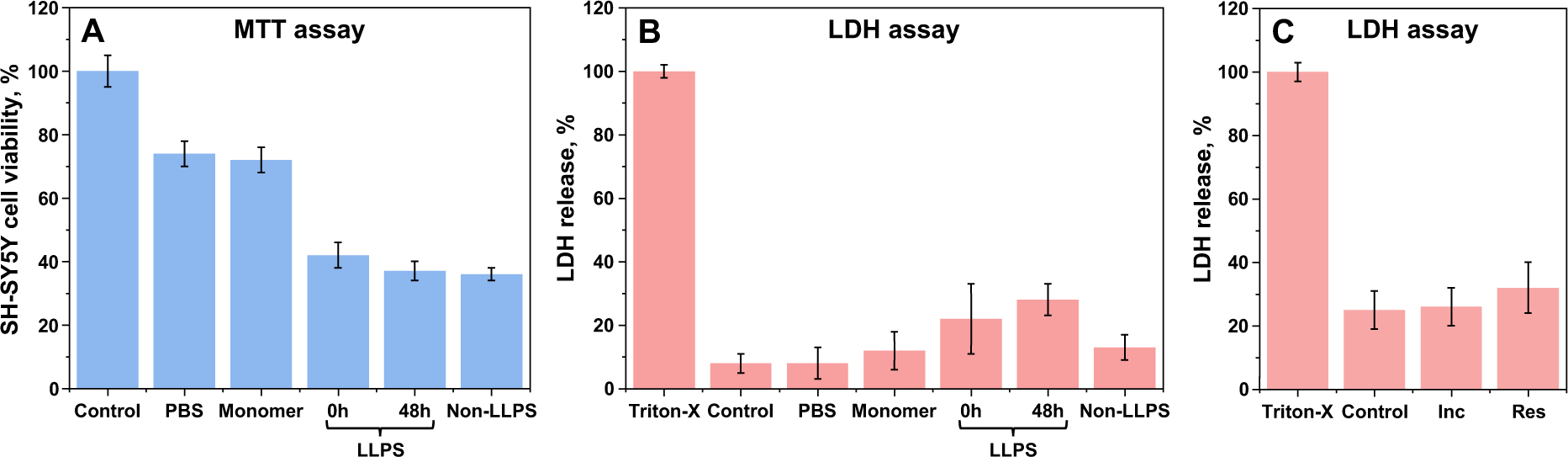
LLPS and non-LLPS aggregate effect on SH-SY5Y human neuroblastoma cells. Initial (0h) and cold-incubated LLPS (48h), as well as non-LLPS aggregate effect on cell metabolic activity (A, MTT assay) and their cytotoxicity (B, LDH assay). LDH assay with LLPS aggregates, after they were incubated in Advanced DMEM medium (Inc) or resuspended into the medium (Res) prior to the assay (C). All initial protein and aggregate concentrations were set to 20 µM prior to the assays (in case of LLPS and non-LLPS aggregate samples, the concentration was based on α-syn in its aggregated state). For each condition, three independent measurements were performed, error bars are for one standard deviation.

The high level of result deviation in the case of 0h incubation LLPS aggregates could be related to the aggregate instability during the LDH assay. To examine this, the aggregates were resuspended into Advanced DMEM and incubated at 37°C for 48 hours. For a control sample, the aggregates were incubated in PBS at 37°C for 48 hours resuspended into the medium prior to the LDH assay. In both cases, there were no significant differences from the control (Figure 5C), suggesting that the predominantly small amorphous aggregates had no notable cytotoxic effect.

## Discussion

The influence of LLPS on amyloidogenic protein aggregation is a new phenomenon that likely plays a role in many different diseases. Here we develop a new method and apply it to isolate amyloid fibrils from LLPS-induced aggregate mixtures. Using this method, we demonstrate the influence of droplet formation on the resulting aggregate structural and morphological variability, self-replication potential and cytotoxicity.

The initial LLPS-induced aggregate sample was composed of a massive quantity of round and amorphous structures with rare instances of amyloid fibrils. In the context of LLPS *in vivo*, the rapid appearance of such an abundance of protein aggregates, both amorphous and fibrillar, can play a significant role in the emergence and/or development of amyloid disorders. The formed structures could either possess cytotoxic properties by themselves ^53,54^ or serve as a catalyst for surface-mediated nucleation for the generation of amyloid nuclei ^55^. During *in vitro* experimental procedures, these meta-stable aggregates also complicate structural analysis of the LLPS-induced structures. In this work, it was discovered that the highly stable amyloid fibrils can be relatively easily isolated from the amorphous structures by the removal of the crowding agent and incubation of the sample under cold conditions. This process of cold-induced dissociation is based on previous observations, where low temperatures were associated with the disassembly of certain types of alpha-synuclein aggregates ^41^.

During the process of cold-incubation and amorphous aggregate dissociation, two interesting factors were observed. The first notable aspect was the incomplete dissolution of non-fibrillar structures, which were visible in samples even after the additional reseeding procedure. This observation, in addition with the gradual reduction in sample optical density upon increasing denaturant concentration, suggests that the LLPS-induced aggregate samples contain a mixture of structures with distinct stabilities against environmental factors. The second interesting factor was the influence of cold conditions on non-LLPS induced aggregates. Based on the FTIR spectra peak-fitting, there was a reduction in stronger hydrogen bonding in the beta-sheet structure, accompanied by an equal increase in weaker hydrogen bonding. Part of the aggregates also dissolved, as seen from the concentration of α-syn in its aggregated state before and after incubation. This effect may also influence part of the LLPS-induced aggregates, by altering their secondary structure or entirely dissolving them.

Another noteworthy observation made during this work was the variety of fibrils that formed via the LLPS pathway. When α-syn was aggregated under conditions which prevent LLPS (no crowding agent and constant agitation), the final fibril structures pertained highly similar morphologies. Oppositely, LLPS conditions resulted in at least five significantly distinct aggregates, which had unique periodicity patterns, average cross-sectional height values and widths. Based on previous reports, it is known that environmental conditions, such as protein concentration, solution ionic strength, pH, temperature and agitation, can influence the resulting fibril structure for both α-syn, as well as other amyloidogenic proteins ^32,33,56,57^. In this work, the initial protein concentration, solution salt concentration, pH and incubation temperature were identical, with the exception of the crowding agent and agitation. This suggests that the protein droplet formation is the main governing factor affecting the resulting aggregate structural variability.

The aforementioned structural variability may stem from two factors: surface-mediated nucleation, which is caused by the abundance of amorphous structures or a concentration-dependent polymorphism due to the high level of protein packing within the droplet. Since the large increase in ThT fluorescence intensity is only observed after 24 hours of incubation, while α-syn droplets display the formation of aggregates before this time-point, it is possible that the amorphous structures act as a surface for α-syn nucleation. The extremely high protein concentration within the droplet may also play a role in the generation of several different fibril types, as we have previously observed that α-syn concentration is directly responsible for fibril structure variability ^33^.

During the LDH assay, it was observed that the stable fibrils within the LLPS-induced aggregate mixtures had the highest cytotoxicity out of all the samples, including fibril formed during non-LLPS conditions. These results suggest that conditions, which promote protein phase separation *in vitro*, may be a step towards better mimicking *in vivo* amyloid fibril formation. As a future perspective, the wide variety of aggregates formed during α-syn LLPS could be examined on a molecular level, using solid-state NMR ^58^ or Cryo-EM ^59^ techniques. The detailed structural analysis of these higher toxicity fibrils could serve in the search for potential anti-amyloid compounds or disease treatment modalities.

