## Supplementary Material for "Liquid-liquid Phase Separation of Alpha-synuclein Increases the Structural Variability of Fibrils Formed during Amyloid Aggregation"

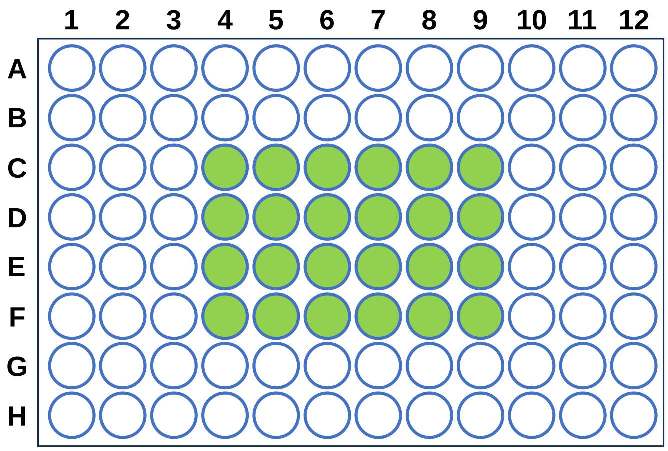


**Figure S1.** Sample placement in 96-well plates during LLPS and aggregation reactions.

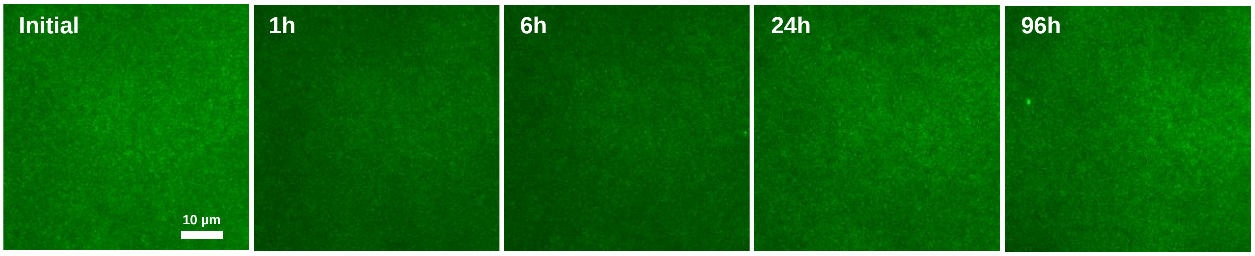


**Figure S2.** Fluorescence microscopy images of eGFP-tagged α-syn samples, incubated under quiescent conditions without PEG. Samples were additionally supplemented with 100 μM ThT to detect amyloid aggregate formation.

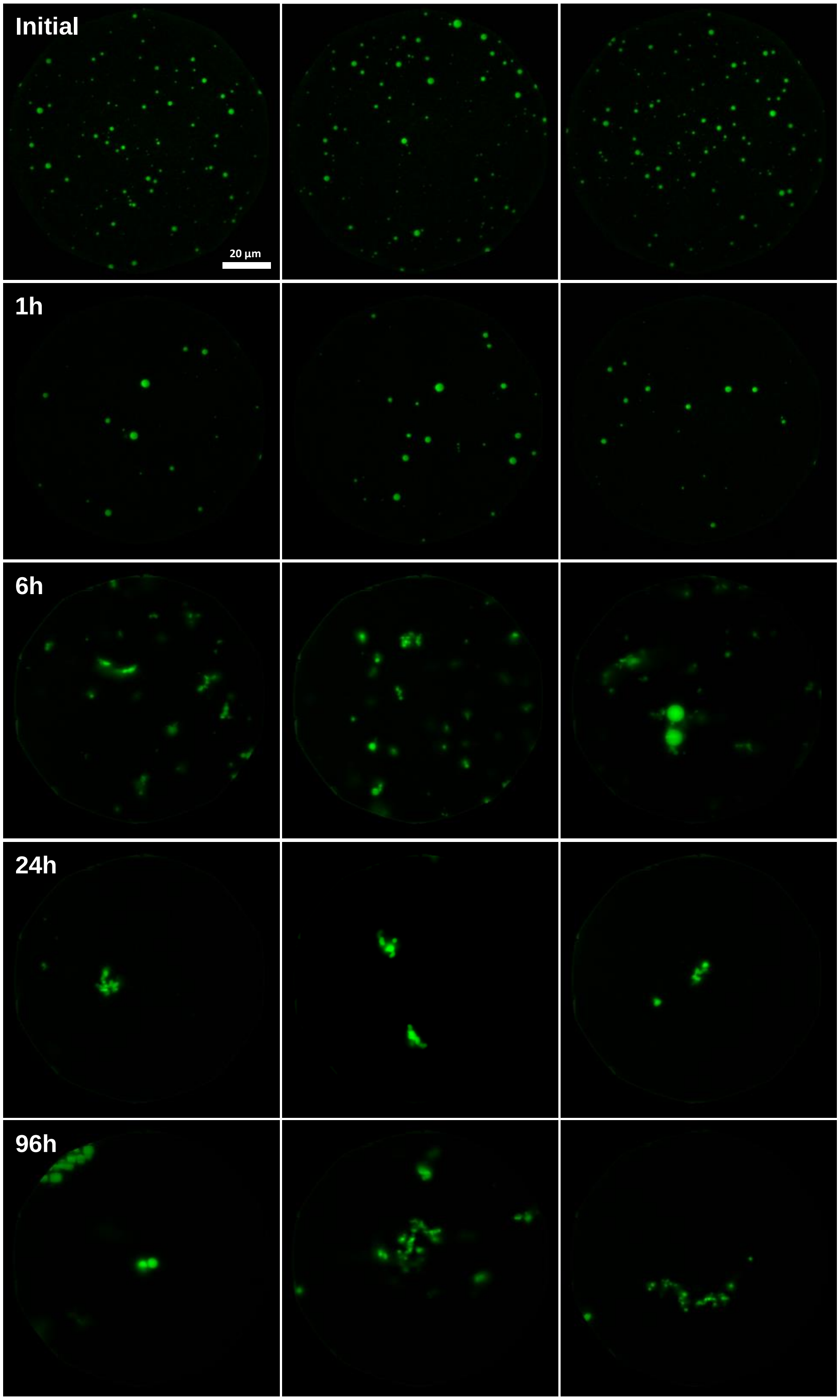


**Figure S3.** Fluorescence microscopy images of eGFP-tagged α-syn samples, incubated under quiescent conditions with 20% PEG.

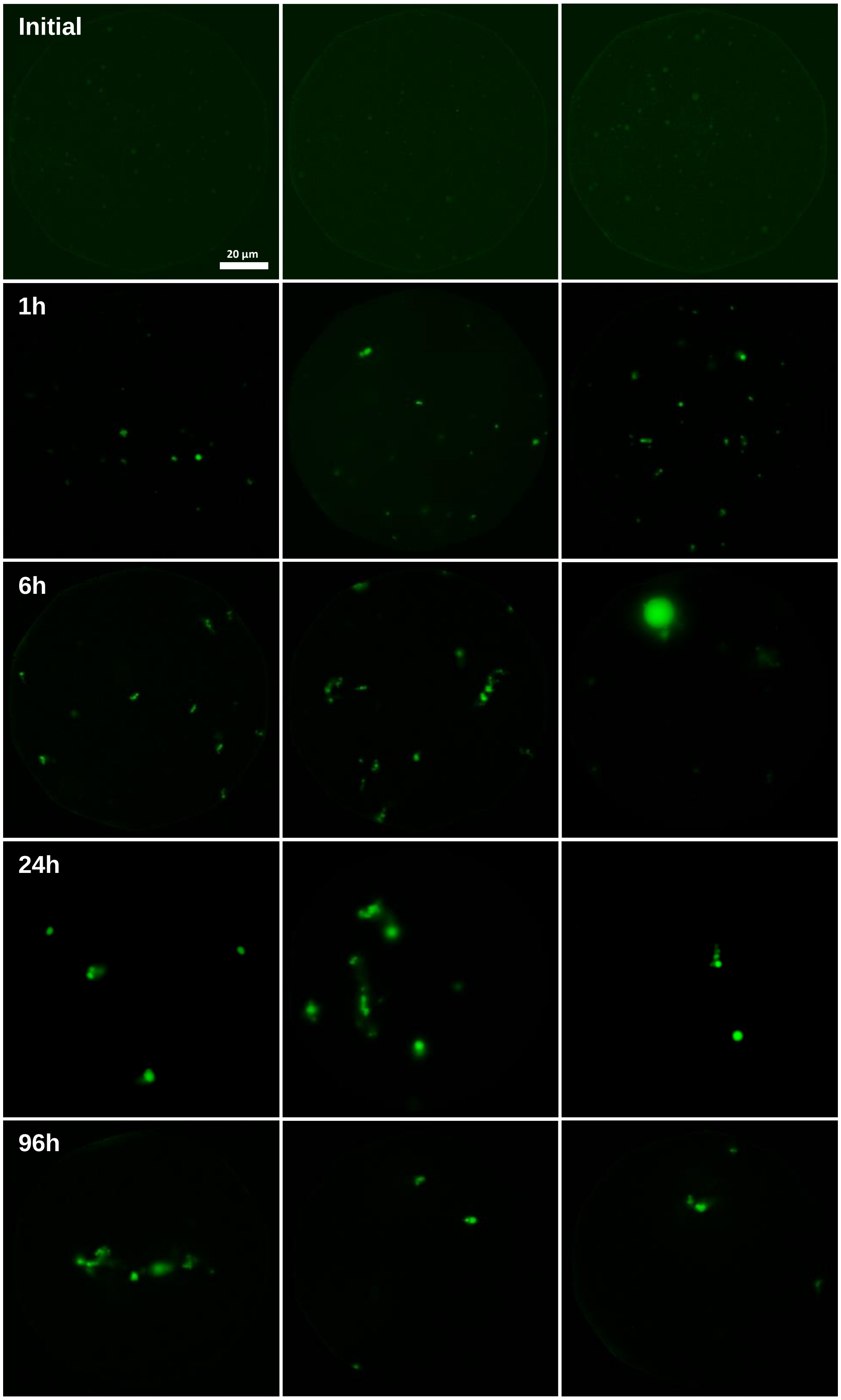


**Figure S4.** Fluorescence microscopy images of α-syn samples, incubated under quiescent conditions with 20% PEG and 100 μM ThT.

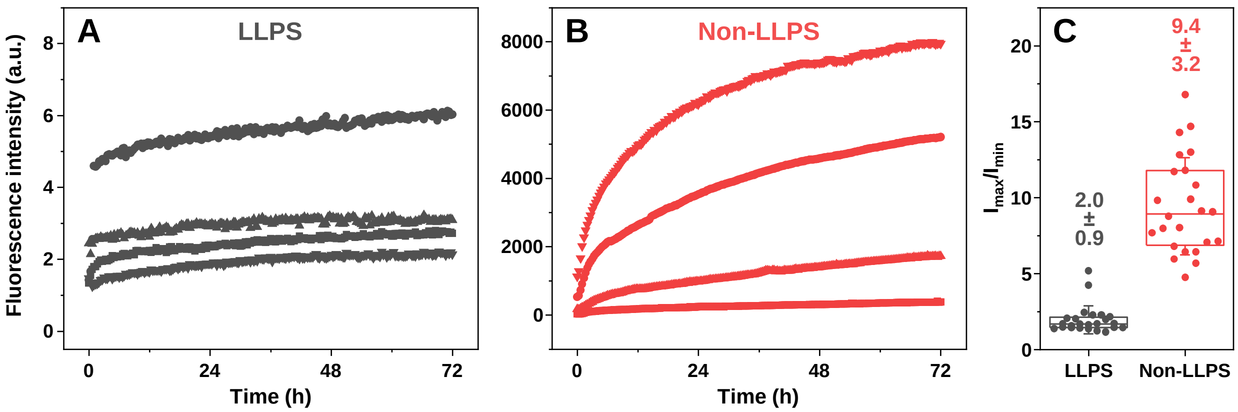


**Figure S5.** Self-replication kinetics of LLPS and non-LLPS aggregates. Example curves of LLPS (A) and non-LLPS (B) self-replication. Ratio between the minimum and maximum fluorescence intensity of all LLPS and non-LLPS samples (C). Error bars are for one standard deviation, box plots represent the interquartile range.

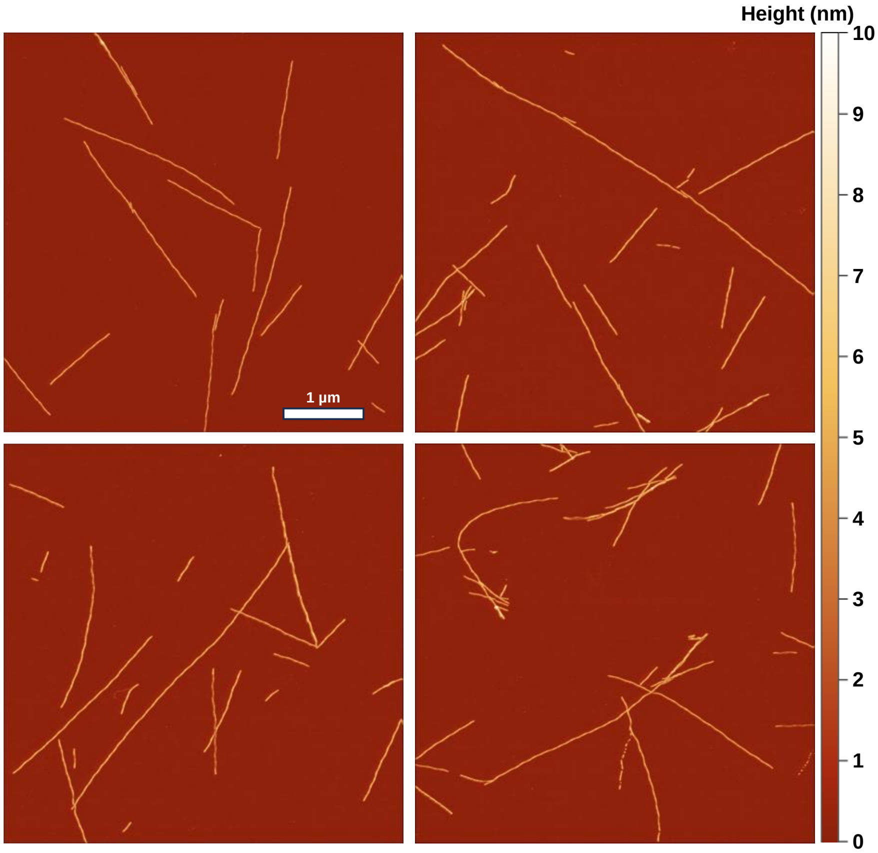


**Figure S6.** Atomic force microscopy images of non-LLPS condition fibrils.

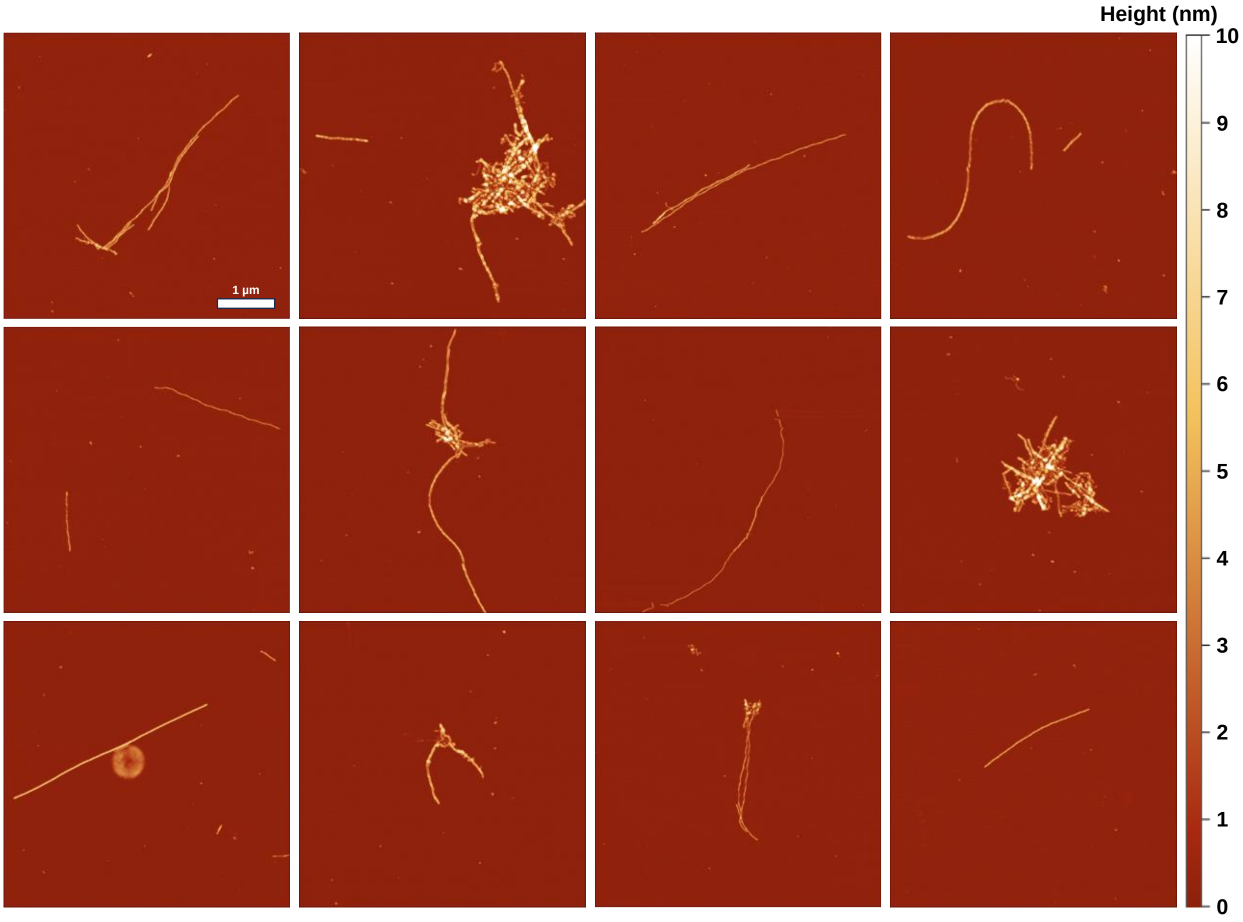


**Figure S7**. Atomic force microscopy images of LLPS condition fibrils.
